# Multi-platforms approach for plasma proteomics: complementarity of Olink PEA technology to mass spectrometry-based protein profiling

**DOI:** 10.1101/2020.08.04.236356

**Authors:** Agnese Petrera, Christine von Toerne, Jennifer Behler, Cornelia Huth, Barbara Thorand, Anne Hilgendorff, Stefanie M. Hauck

## Abstract

The plasma proteome is the ultimate target for biomarker discovery. It stores an endless amount of information on the pathophysiological status of a living organism, which is however still difficult to comprehensively access. The high structural complexity of the plasma proteome can be addressed by either a system-wide and unbiased tool such as mass spectrometry (LC-MS/MS) or a highly sensitive targeted immunoassay such as the Proximity Extension Assays (PEA). In order to address relevant differences and important shared characteristics, we tested the performance of LC-MS/MS in data-dependent and -independent acquisition modes and PEA Olink to measure circulating plasma proteins in 173 human plasma samples from a Southern German population-based cohort. We demonstrated the measurement of more than 300 proteins with both LC-MS/MS approaches applied, mainly including high abundance plasma proteins. By the use of the PEA technology, we measured 728 plasma proteins, covering a broad dynamic range with high sensitivity down to pg/ml concentrations. In a next step, we quantified 35 overlapping proteins with all three analytical platforms, verifying the reproducibility of data distributions, measurement correlation and gender-based differential expression. Our work highlights the limitations and the advantages of both, targeted and untargeted approaches, and prove their complementary strengths. We demonstrated a significant gain in proteome coverage depth and subsequent biological insight by platforms combination – a promising approach for future biomarker and mechanistic studies.

## Introduction

The advent of omics technologies gave rise to the era of the precision medicine. The concept that health care can be personalized according to our genes, lifestyle and environment is becoming more concrete and its implementation in the clinical practice is a tangible reality (1). Genes, metabolites and protein landscapes are nowadays simultaneously explored using a multitude of state-of-the-art technologies, enabling us to capture the whole picture of a biological system (2). The contribution of omics technologies to personalized therapeutic approaches is centered around biomarkers, the essence of precision medicine (3). The tailored management of a disease relies intrinsically on a biomarker-driven approach, hence the urge to develop novel biomarker discovery platforms excelling in sensitivity, specificity and multiplexing. The largest proportion of laboratory clinical tests is based on proteins circulating in blood plasma, highlighting the importance of the plasma proteome in clinical diagnostics (4). Plasma represents the elected body fluid for biomarker discovery among others because of the excellent accessibility and the rich protein repertoire, which mirrors the pathophysiological status of a living organism. However, the outstanding dynamic range of plasma proteins, which covers more than 10 orders of magnitude, places substantial limits for mass spectrometry (MS) based proteomic platforms (5). Meeting these limitations, MS-based plasma proteome technologies have dramatically improved, and recently proposed workflows based on the combination of multiple biomarkers with clinical metadata, hold great promise to discover disease specific-protein patterns (6–8). However, pre-analytical variability, depth of proteome coverage and robustness of discovered biomarkers still remain the hardest challenges for the MS field (9). MS data-dependent acquisition methods (DDA) with improved MS1 level performance (termed BoxCar) enabled to identify 580 human plasma proteins in at least 25 % of the samples in a longitudinal cohort (10). With the same approach and similar proteome coverage, Niu et al. identified a panel of six new biomarker candidates for non-alcoholic fatty liver disease (11). Increasingly, large-scale plasma MS studies are performed using a data-independent acquisition (DIA) technology. This approach does not rely on precursor intensity for fragmentation but rather fragments in a systematic and unbiased fashion all precursors in specified mass ranges, therefore providing an opportunity to improve plasma-based biomarker discovery studies. Bruderer et al. conducted the so far largest human plasma DIA study using a capillary flow DIA setup (12); the high throughput and robust workflow enabled profiling of 565 proteins with 74% data set completeness in 1508 plasma samples. Targeted methods based on immuno-detection of specific proteins are receiving increasing attention in plasma proteomics (13). The proximity extension assay (PEA) is an emerging affinity-based proteomic technology developed by Olink Proteomics (Uppsala, Sweden), which merges quantitative real-time PCR with multiplex immunoassays. The basis of PEA is a dual-recognition of a targeted biomarker through a matched pair of antibodies labelled with unique DNA oligonucleotides. Here, biomarker-specific DNA “barcodes” are quantified by microfluidic qPCR allowing for high throughput relative quantification of up to 1161 human plasma proteins by using a few microliters of bio fluid (1 μl for the quantification of 92 biomarkers). The currently available 14 human panels, each comprising of 92 biomarkers, are designed to cover a dynamic range over 10 orders of magnitude, providing accurate quantification below picogram per milliliter (pg/ml) levels. The impact of the PEA targeted proteomic platform in the human plasma biomarker field is undeniable: the last five years saw an exponential increase of research articles presenting clinical data measured using the PEA technology. More than 230 original publications report PEA-based biomarker research on human plasma, overgrowing the annual number of articles presenting MS-based approaches. Previous studies have shown positive correlation of PEA with SOMAscan data in multi-platform affinity studies (14, 15), as well as complementarity of mass spectrometry to SOMAscan, an aptamer-based targeted proteomics platform, using stem cell comparisons as a model (16). Here we aim to benchmark PEA technology against nanoliquid chromatography (LC) MS approach in DDA and DIA mode using 173 blood plasma samples from the population-based KORA (Cooperative Health Research in the Region of Augsburg) study. We characterized and compared the performances of these analytical platforms to explore the plasma proteome. Our multi-platform proteomics approach shows the complementarity of PEA and untargeted LC-MS/MS in the identification and quantification of plasma proteins of clinically relevant samples, while investigating potentials and limitations of each technology.

## Experimental Procedures

### Samples

The plasma samples were collected from participants of the population-based KORA F4 study. The KORA F4 study was carried out in 2006–2008 and included 3080 participants. Investigations were carried out in accordance with the Declaration of Helsinki, and written informed consent was obtained from all participants. The study was approved by the Ethics committee of the Bavarian Chamber of Physicians (17). Plasma samples from 90 men and 83 women KORA F4 participants within the age range of 62–74 years were randomly selected from a subset of KORA F4 participants with deep phenotyping data (18–20).

### Sample preparation for mass spectrometry

Plasma samples were prepared using PreOmics’ iST Kit (Preomics GmbH, Martinsried, Germany) according to manufacturer’s specifications. After drying, the peptides were resuspended in 3% ACN and 0.5% TFA acid. The HRM Calibration Kit (Biognosys, Schlieren, Switzerland) was added to all of the samples according to manufacturer’s instructions.

### Mass spectrometry measurements

MS data were acquired in data-independent acquisition (DIA) mode as well as data-dependent acquisition (DDA) mode on a Q Exactive (QE) high field (HF) mass spectrometer (Thermo Fisher Scientific Inc.). Approximately 0.5 μg per sample were automatically loaded to the online coupled RSLC (Ultimate 3000, Thermo Fisher Scientific Inc.) HPLC system. A nano trap column was used (300 μm inner diameter (ID) × 5 mm, packed with Acclaim PepMap100 C18, 5 μm, 100 Å; LC Packings, Sunnyvale, CA) before separation by reversed phase chromatography (Acquity UPLC M-Class HSS T3 Column 75μm ID x 250mm, 1.8μm; Waters, Eschborn, Germany) at 40°C. For DIA acquisition, peptides were eluted from column at 250 nl/min using increasing acetonitrile (ACN) concentration (in 0.1% formic acid) from 3% to 40 % over a 45 minutes gradient. The HRM DIA method consisted of a survey scan from 300 to 1500 m/z at 120 000 resolution and an automatic gain control (AGC) target of 3e6 or 120 ms maximum injection time (IT). Fragmentation was performed via higher energy collisional dissociation (HCD) with a target value of 3e6 ions determined with predictive AGC. Precursor peptides were isolated with 17 variable windows spanning from 300 to 1500 m/z at 30 000 resolution with an AGC target of 3e6 and automatic injection time. The normalized collision energy (CE) was 28 and the spectra were recorded in profile type. For DDA acquisition mode, peptides were eluted as for DIA acquisition from 3% to 40 % over a 95 minutes gradient. The MS spectrum was acquired with a mass range from 300 to 1500 m/z at resolution 60 000 with AGC set to 3 x 10^6^ and a maximum of 50 ms IT. From the MS prescan, the 10 most abundant peptide ions were selected for fragmentation (MS/MS) if at least doubly charged, with a dynamic exclusion of 30 seconds. MS/MS spectra were recorded at 15 000 resolution with AGC set to 1 x 10^5^ and a maximum of 100 ms IT. CE was set to 28 and all spectra were recorded in profile type. SRM-MS measurement of MBL2 is described by Von Toerne et al. (21).

### Generation of the spectral library for DIA data analysis

For the spectral library an in-house generated plasma library was merged with publically available LS-MS/MS runs. The in-house library encompassed 89 files of plasma and serum preparations, spiked with the HRM Calibration Kit, were analyzed using Proteome Discoverer (Version 2.1, ThermoFisher Scientific) using the search engine node Byonic (Version 2.0, Proteinmetrics, San Carlos, CA). Peptide and protein FDRs were filtered to satisfy the 1% peptide level. The peptide spectral library was generated in Spectronaut (Biognosys, Schlieren, Switzerland) with default settings using the Proteome Discoverer result file. Spectronaut was equipped with the Swissprot human database (Release 2017.02, 20194 sequences, www.uniprot.org) with a few spiked proteins (e.g., Biognosys iRT peptide sequences). The second part of the library was generated from 95 publically available runs of fractionated plasma from (10). The spectral library was built in Spectronaut 12 - Pulsar directly, applying peptide and protein FDR of 1%. Both databases were merged in Spectronaut 12 according manufacturer settings. The final spectral library generated in Spectronaut contained 1838 protein groups and 37079 peptide precursors.

### Spectronaut analysis and data processing of DIA measurements

The DIA-MS data were analyzed using the Spectronaut software (version 12; Biognosys, Schlieren, Schwitzerland) as described previously (22). Due to the explorative nature of the study, no protein FDR on the identification level was applied. Quantification is based on the summed up abundances of proteotypic peptides with data filtering set to Qvalue required. Abundance values were normalized based on retention time dependent local regression model (23).

### Proteome Discoverer analysis and data processing of DDA measurements

Proteome Discoverer 2.3 software (Thermo Fisher Scientific; version 2.3.0.523) was used for peptide and protein identification via a database search (Sequest HT search engine) against Swissprot human database (release 2017_2, 20237 sequences), considering full tryptic specificity, allowing for up to two missed tryptic cleavage sites, precursor mass tolerance 10 ppm, fragment mass tolerance 0.02 Da. Carbamidomethylation of Cys was set as a static modification. Dynamic modifications included deamidation of Asn and Gln, oxidation Met; and a combination of Met loss with acetylation on protein *N*-terminus. Percolator (24) was used for validating peptide spectrum matches and peptides, accepting only the top-scoring hit for each spectrum, and satisfying the cutoff values for FDR <1%, and posterior error probability <0.01. The final list of proteins complied with the strict parsimony principle. The quantification of proteins, after precursor recalibration, was based on abundance values (area under curve) for unique peptides. Abundance values were normalized in a retention time dependent manner. The protein abundances were calculated summing the abundance values for admissible peptides.

### Olink PEA measurement

The relative abundance of 736 protein biomarkers in plasma was determined using the Olink C-MET, CVDII, CVDIII, ONCII, ONCIII, DEV, IMM and NEU panels. Each panel of 92 proteins was screened using 1 μl of plasma. Protein abundance was quantified by real-time PCR using the Fluidigm BioMark™ HD real-time PCR platform (25). Briefly, each protein is targeted by a specific pair of oligonucleotide-labeled antibodies and when the two oligonucleotides are placed in close proximity, a DNA polymerization event generates a specific PCR product which is detected and quantified using real-time PCR. For each proximity extension assay (PEA) measurement, a limit of detection (LOD) is calculated based on negative controls included in each run and measurements below this limit were removed from further analysis. The Olink PEA data are reported in NPX values (normalized protein expression levels) which is on log2-scale, calculated as previously described (25). To avoid batch effects, samples were randomized across plates. Each plate included inter-plate controls which were used to adjust for any plate difference. NPX values were intensity normalized with the plate median for each assay as the normalization factor (Intensity Normalization v.2). Samples and proteins that did not pass the quality control were excluded. For assays measured in multiple panels, only one measurement was included in the analysis (AREG-OID00728, IL6-OID00390, SCF-OID00408); the following assays have been therefore removed: OID00947, OID00666, OID01009, OID00684. A complete list of the measurements is included in Supplemental Table S2. Details about PEA technology, assays performance and validation data are available from the manufacturer (www.olink.com).

### Experimental Design and Statistical Rationale

173 human plasma samples from the Southern Germany population-based cohort KORA were used for this study. An in-house plasma pool standard was included in MS and PEA measurements to determine the intra- and inter-assay coefficient of variance. Negative and positive controls for Olink PEA experiments are provided for each measurement by the manufacturer. All statistical analyses were performed in R version 3.6.2 (https://www.r-project.org/) and RStudio (https://rstudio.com/) version 1.2.5001. PEA data was input into R using the OlinkAnalyze package, freely available at https://github.com/Olink-Proteomics/OlinkRPackage. Before statistical analysis, the MS data was log2 transformed.

### Bioinformatics analysis

For protein enrichment analysis, the protein IDs subsets were submitted to ShinyGO v0.61 (http://bioinformatics.sdstate.edu/go/), setting p-value cutoff to 0.05. For generation of the network of proteins, we used STRING (version 11.0, https://string-db.org). STRING was employed using “medium confidence” regarding the interaction score, with a score limit of 0.4. The thickness of the connection lines indicates the strength of the data support. Venn diagram was generated using FunRich (http://www.funrich.org/).

## Results

### 1. Qualitative comparative analysis of the PEA and LC-MS/MS datasets

Our benchmarking study was conducted on 173 plasma samples from the longitudinal population-based KORA (Cooperative Health Research in the Region of Augsburg) study. Eight Olink panels were selected to cover mostly proteins with higher expected abundances in plasma and hence more potential overlap with LC-MS/MS-based identifications. By using PEA, we screened 736 protein biomarkers (Fig.1A). Three proteins were measured in multiple panels (SCF, IL6 and AREG) and measurements across panels showed a strong positive correlation with Spearman R coefficient ranging from 0.84 to 0.95 (**Supplemental Table S1**). After quality control and normalization (complete list of PEA results in **Supplemental Table S2**), 728 proteins were detected in more than 25% of the samples and more than 90% of the targeted proteins (666) were detected above the limit of detection in all samples. Mass spectrometrybased measurements of undepleted plasma samples were performed in DDA and DIA modes. By using a standard DDA-MS workflow a total of 368 proteins (3042 peptides) were identified and quantified cumulatively in all samples (protein FDR < 0.05, at least one unique peptide/protein), of which 329 were directly measured (not based on match-between run identification) in more than 25% of the samples (protein and peptide lists available in **Supplemental Table S3**-**S4**). For DIA-MS measurement, we generated a comprehensive spectral library from human undepleted plasma and serum consisting of 1838 proteins (37079 peptide precursors). Using our spectral library, we quantified a total of 734 proteins in the plasma samples (12314 peptides), an extended proteome coverage which narrowed to 379 proteins when filtering the direct measurements in more than 25% of the samples. This observation can be attributed to our stringent peptide threshold q-value which was set to ≤ 0.01 (protein and peptide lists available in **Supplemental Table S5-S6**). Also for the MS platform, we assessed the intra-assay reproducibility by correlating the repeated measurements (n=12) of a plasma pool. For both DDA- and DIA-MS, we observed an excellent intra-platform correlation (Spearman R= 0.99 and R=0.98 for DDA- and DIA-MS, respectively).

**FIGURE 1.**
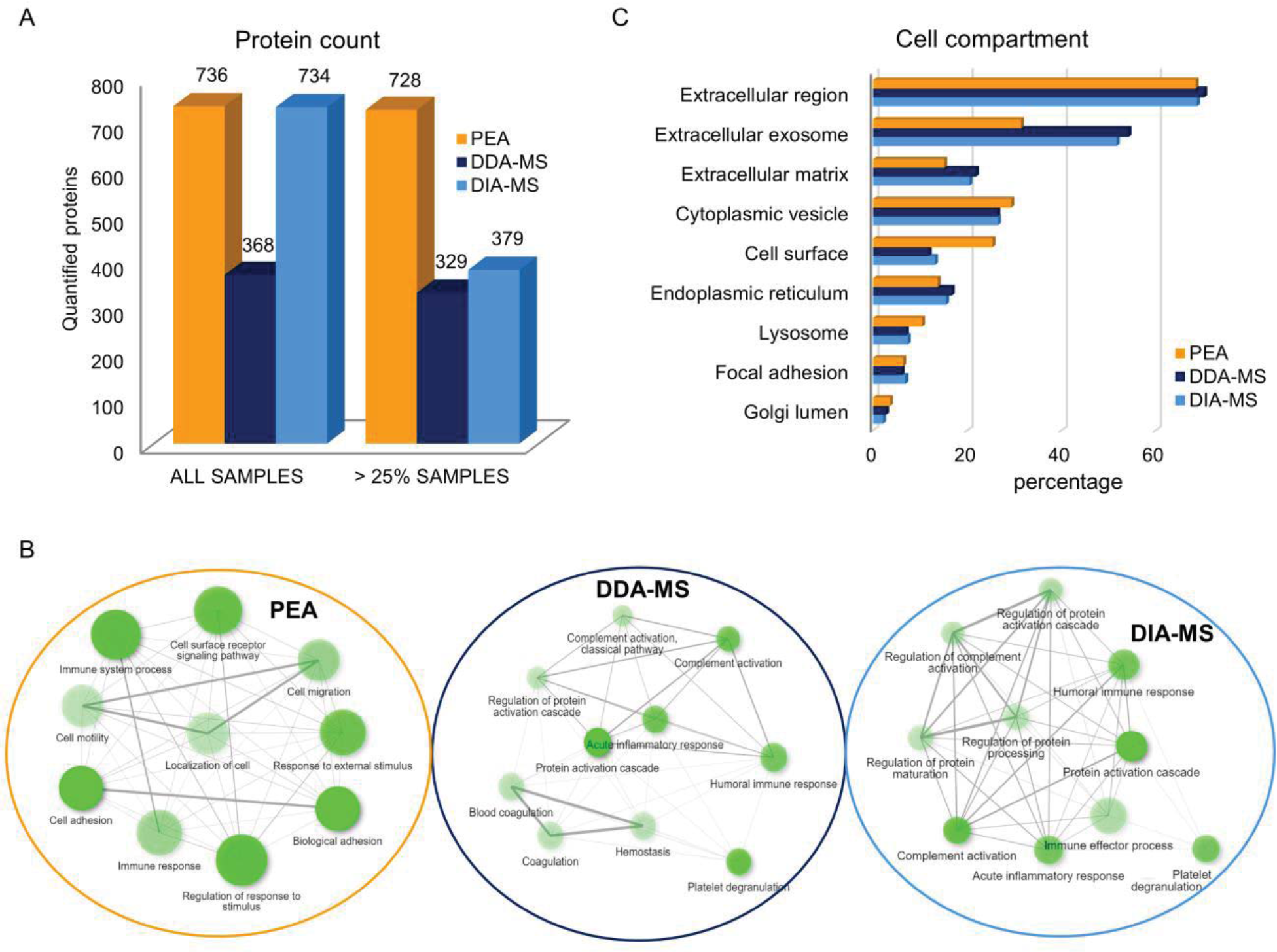
A. Number of proteins identified and quantified by PEA, DDA- and DIA-MS, cumulatively in all samples or in more than 25% of the samples. B. Ten most significant GO biological processes in the subproteomes identified by PEA, DDA-MS and DIA-MS. Darker nodes are more significantly enriched gene sets. Bigger nodes represent larger gene sets. Thicker edges represent more overlapped genes. C. Enrichment analysis based on GO cell compartment for each platform.

We next characterized the subproteome targeted by PEA, DDA-MS and DIA-MS (Fig.1B). We used the GO classification to explore the biological processes represented by proteins identified in more than 25% of the samples in each analytical platform. Among top enriched biological processes in the PEA targeted subproteome emerged signaling pathways, immune system and biological adhesion (FDRs 4.98E^−89^, 9.05E^−78^ and 2.48E^−108^, respectively); specifically, proteins involved in cell surface receptor signaling pathway represented more than 40% of the screened proteins. Among them, we measured 37 CD (cluster of differentiation) markers and 12 TNF (Tumor necrosis factor) Receptor Superfamily Members. The high sensitivity of the PEA technology allowed the measurement of low abundance proteins such interleukins and chemokines (for a total of 25 different molecules), peptide hormones (5) and growth factors including TFG-alpha (Transforming growth factor alpha, P01135) (25). Given their extremely low abundance at steady state, these low-molecular weight signaling proteins are hardly detected by MS. We indeed measured only two chemokines by MS (Platelet basic protein, P02775 and Platelet factor 4, P02776). The most represented biological process in the DDA- and DIA-MS datasets was immune response, which included 33% and 34% of the total IDs for DDA-MS and DIA-MS (FDRs 3E^−24^ and 1E^−31^, respectively). A large number of immunoglobulins (many of which are variable domains of immunoglobulins for heavy and light chains) and complement factors were detected by MS (107 and 101 protein IDs for DIA-MS and DDA-MS, respectively). The 14 available PEA panels do not include immunoglobulins, but instead 26 immunoglobulin receptors and immunoglobulin domain-containing proteins. Unlike the PEA-targeted subproteome, apolipoproteins were highly represented in the MS datasets. Indeed, only APOM (Apolipoprotein M, O95445) was targeted by PEA, whereas 14 different apolipoproteins (including APOM) were identified by both DDA- and DIA-MS workflows. As shown in Fig.1C, nearly 70% of the proteins identified by all three platforms are located in the extracellular region, which relates very well with the nature of the body fluid examined here (FDRs < E^−88^). The high representation of extracellular exosome-located proteins in the MS subproteomes reflects the large abundance of immunoglobulins, coagulation factors and complement proteins which are known to be associated to extracellular vesicles (FDRs < E^−91^) (26). On the other side, the doubled proportion of cell surface proteins identified by PEA compared to MS mirrors the overrepresentation of cell signaling molecules targeted by this approach (FDR 7.4E^−90^).

### 2. Overlap analysis of proteins identified by PEA, DDA- and DIA-MS

To allow for a robust and reliable comparison of proteins identified by all three analytical platforms, we overlapped the proteins detected in more than 25% of the samples. As shown in Fig. 2A, over 60% of the total proteins detected by MS were quantified in both DDA and DIA modes (276 IDs). A far narrower overlap of 52 proteins resulted from a comparison of proteins measured by both MS and PEA. Most interestingly, 35 proteins targeted by PEA were also measured by MS in both DDA and DIA modes (Table 1). As expected, the overlapping proteins between PEA and MS are mainly covered by PEA panels enriched for high abundant proteins, which require pre-dilution of plasma, such as C-MET and CVD III (23 and 7 proteins, respectively). We also find an overlap of 5 proteins within other panels which apply undiluted plasma (Fig. 2A). However, these five proteins are also very abundant in human plasma, as validated by Olink Proteomics during measurement of biomarker analytical concentration ranges (see Validation Data Documents, https://www.olink.com/resources-support/document-download-center/). STRING analysis of the overlapping proteins highlighted a cluster of proteins involved in the complement/activation cascades, as shown in Fig.2B. The majority of the proteins are located in extracellular region, except for the four cell membrane proteins CDH5 (Cadherin-5, P33151), GP1BA (Platelet glycoprotein Ib alpha chain, P07359), ICAM2 (Intercellular adhesion molecule 2, P13598) and LYVE1 (Lymphatic vessel endothelial hyaluronic acid receptor 1, Q9Y5Y7). Outside of the overlap, it is interesting to notice the proteins targeted uniquely by PEA or both MS methods. Although over 600 proteins targeted by PEA are not detected by MS, it is still noteworthy that 380 plasma proteins can be uniquely explored by MS when combining both acquisition methods. Finally, complementing all three platforms, we reached a proteome coverage of 1104 proteins which are reliably identified and quantified in human plasma. In order to verify the complementarity of the platforms, statistical significance (and consequently biological insights) needs to be replicated. We therefore looked at platform reproducibility of the 35 overlapping proteins across methods based on three main components: data distribution, correlation and variable-based differential expression.

**FIGURE 2.**
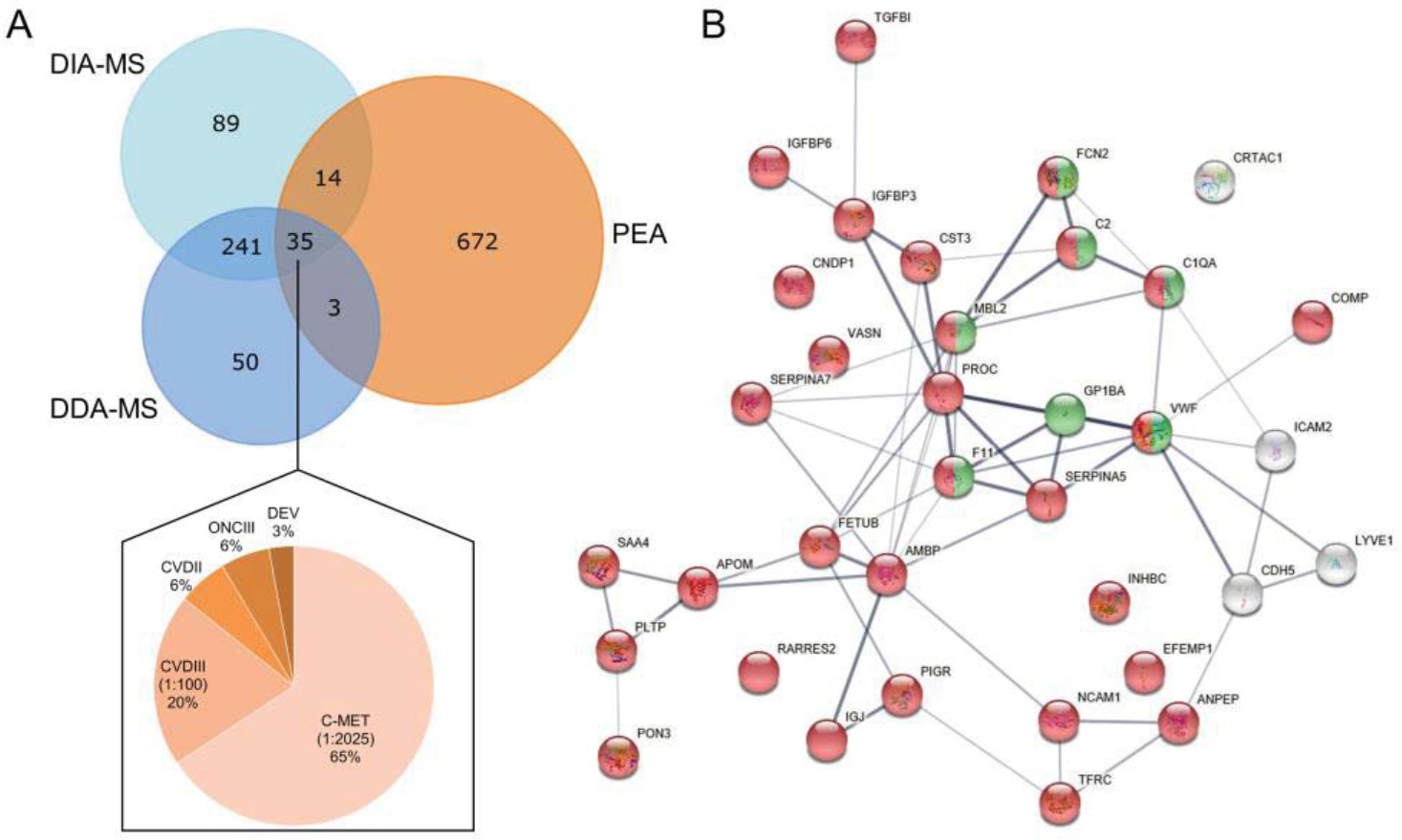
Comparison between proteins found in >25% samples by PEA, DDA- and DIA-MS. A. Venn diagram of proteins detected in more than 25% of the samples by PEA, DDA- and DIAMS. For assays measured in multiple Olink panels, only one measurement was included in the analysis (see Exp. Procedures). Distribution of the 35 overlapping proteins within the eight Olink panels (C-MET: cardiometabolic, which requires a sample dilution 1:2025; CVDIII: cardiovascular III, dilution factor 1:100; CVDII: cardiovascular II; ONCIII: oncology III; DEV: development, no sample dilution is required for the last three panels). B. STRING analysis of the 35 overlapping proteins. Color legend: red – extracellular region (GO:0005576; FDR 5.66e^−20^), green - protein activation cascade (GO:0072376; FDR 9.64e^−08^), grey – background.

**TABLE 1.** Overlapping proteins in PEA and MS measurements. (C-MET: cardiometabolic; CVDII: cardiovascular II; CVDIII: cardiovascular III; ONCIII: oncology III; DEV: development)

**Proteins measured by DDA-MS, DIA-MS and PEA**
| Uniprot ID | Gene name | Protein name | Olink Panel |
| --- | --- | --- | --- |
| Q15582 | TGFBI | Transforming growth factor-beta-induced protein ig-h3 | C-MET |
| P55058 | PLTP | Phospholipid transfer protein | C-MET |
| P05543 | SERPINA7 | Thyroxine-binding globulin | C-MET |
| P17936 | IGFBP3 | Insulin-like growth factor-binding protein 3 | C-MET |
| P05154 | SERPINA5 | Plasma serine protease inhibitor | C-MET |
| P24592 | IGFBP6 | Insulin-like growth factor-binding protein 6 | C-MET |
| P04070 | PROC | Vitamin K-dependent protein C | C-MET |
| P01034 | CST3 | Cystatin-C | C-MET |
| P13591 | NCAM1 | Neural cell adhesion molecule 1 | C-MET |
| P03951 | F11 | Coagulation factor XI | C-MET |
| P49747 | COMP | Cartilage oligomeric matrix protein | C-MET |
| P11226 | MBL2 | Mannose-binding protein C | C-MET |
| Q15485 | FCN2 | Ficolin-2 | C-MET |
| Q9UGM5 | FETUB | Fetuin-B | C-MET |
| Q9NQ79 | CRTAC1 | Cartilage acidic protein 1 | C-MET |
| P35542 | SAA4 | Serum amyloid A-4 protein | C-MET |
| Q96KN2 | CNDP1 | Beta-Ala-His dipeptidase | C-MET |
| Q12805 | EFEMP1 | EGF-containing fibulin-like extracellular matrix protein 1 | C-MET |
| P07359 | GP1BA | Platelet glycoprotein Ib alpha chain | C-MET |
| Q9Y5Y7 | LYVE1 | Lymphatic vessel endothelial hyaluronic acid receptor 1 | C-MET |
| O95445 | APOM | Apolipoprotein M | C-MET |
| Q6EMK4 | VASN | Vasorin | C-MET |
| P06681 | C2 | Complement C2 | C-MET |
| P01833 | PIGR | Polymeric immunoglobulin receptor | CVD II |
| P02760 | AMBP | Protein AMBP | CVD II |
| P33151 | CDH5 | Cadherin-5 | CVD III |
| P02786 | TR | Transferrin receptor protein 1 | CVD III |
| P15144 | ANPEP | Aminopeptidase N | CVD III |
| Q15166 | PON3 | Paraoxonase | CVD III |
| Q99969 | RARRES2 | Retinoic acid receptor responder protein 2 | CVD III |
| P13598 | ICAM2 | Intercellular adhesion molecule 2 | CVD III |
| P04275 | VWF | von Willebrand factor | CVD III |
| P55103 | INHBC | Inhibin beta C chain | DEV |
| P01591 | JCHAIN | Immunoglobulin J chain | ONC III |
| P02745 | C1QA | Complement C1q subcomponent subunit A | ONC III |

#### 2.1 Data distribution of the overlapping proteins identified by PEA, DDA- and DIA-MS

The first port of entry for statistical significance is the distribution of the data. This determines not only the statistical analyses to be applied, but is assumed to represent the underlying population. In order to measure similarity between distributions, we employed the nonparametric two-sample Kolmogorov-Smirnov (KS) test. This was done on z-score transformed (centered and scaled) data for evaluating overall distribution shape. The KS test was set to reject the null hypothesis of equal distribution at an alpha level of 0.05 (no multiple hypothesis testing correction was applied). As shown in figure 3A, the majority of the distributions of the overlapping proteins did not significantly differ between technologies for the z-score transformed data. For instance, 33 and 31 out of 35 proteins were not rejected between PEA and DDA-MS or DIA-MS, respectively. Indeed, the reproducibility of distributions between PEA and MS was as high as within MS platforms. The distributions of some exemplary proteins is shown in Figure 3B and full KS test results are displayed in **Supplemental Table S7**.

**FIGURE 3.**
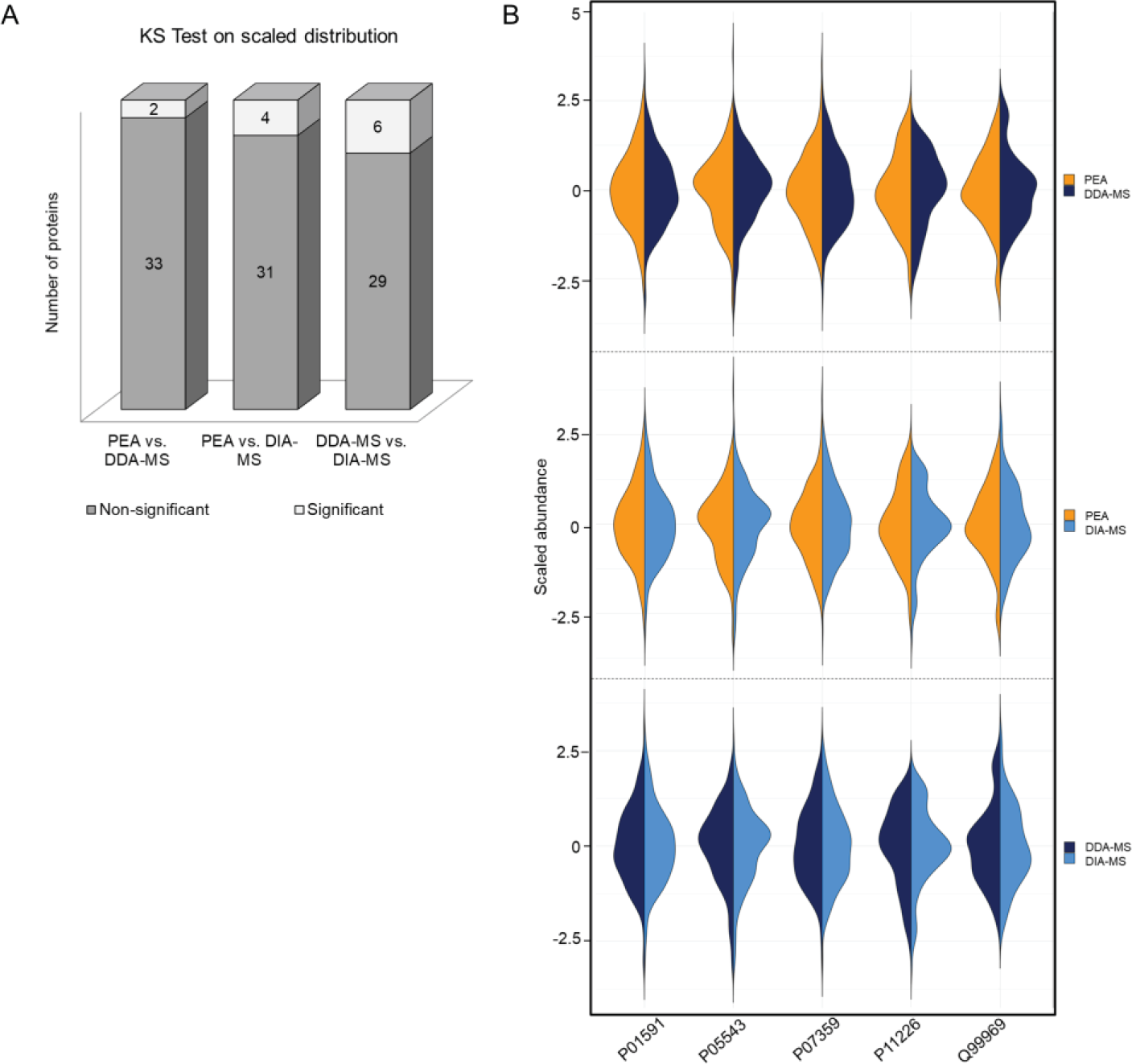
Data distribution of the 35 overlapping proteins. A. Non-parametric two-sample Kolmogorov-Smirnov (KS) test on z-score transformed scaled data. The histogram shows the number of proteins with significant or non-significant difference of the data distribution between platforms. B. Violin plots of the z-score transformed scaled data showing the distribution shape of five exemplary overlapping proteins.

#### 2.2 Spearman correlation of the overlapping proteins identified by PEA, DDA- and DIA-MS

Secondly, Spearman correlation coefficients were calculated for the 35 proteins. As shown in Table 2, MS measurements using DDA and DIA had a fair correlation for the majority of the proteins (23 proteins with DDA-DIA Spearman correlation R > 0.5). PEA measurements showed a slightly higher degree of correlation with DDA-MS than DIA-MS, i.e. respectively 14 and 10 proteins with Spearman R > 0.5. Six proteins had a moderate positive correlation (Spearman R between 0.5 and 0.8) in all three platforms, and two proteins had high positive correlation (Spearman R > 0.8). Among them, only MBL2 (Mannose-binding protein C, P11226) displayed excellent correlation between platforms (Spearman R > 0.9) (Figure 4A). Further investigation showed that for the majority of overlapping proteins the data spread around the mean was quite narrow. To illustrate the spread, a KS test was applied on centered data only. Full KS test results can be found in **Supplemental Table S8**. As expected, a large number of proteins did not fulfill the assumption of equal spread (rejection). As can be seen in Fig. 4B, among the nine proteins with equal spread (no rejection) measured by all three platforms, the large spread of MBL2 is evident. Spread is a prerequisite for correlation calculation, or in other words, there needs to be an inherent direction of the data in both techniques to observe a significant correlation. The only protein fulfilling these criteria is MBL2, which showed excellent correlation. Measurement of plasma levels of MBL2 by selected reaction monitoring-mass spectrometry (SRM-MS) in the cohort used here has been previously published by our group (19, 21). PEA measurements positively correlate with the SRM-MS data with a Spearman coefficient R= 0.84.

**TABLE 2.**
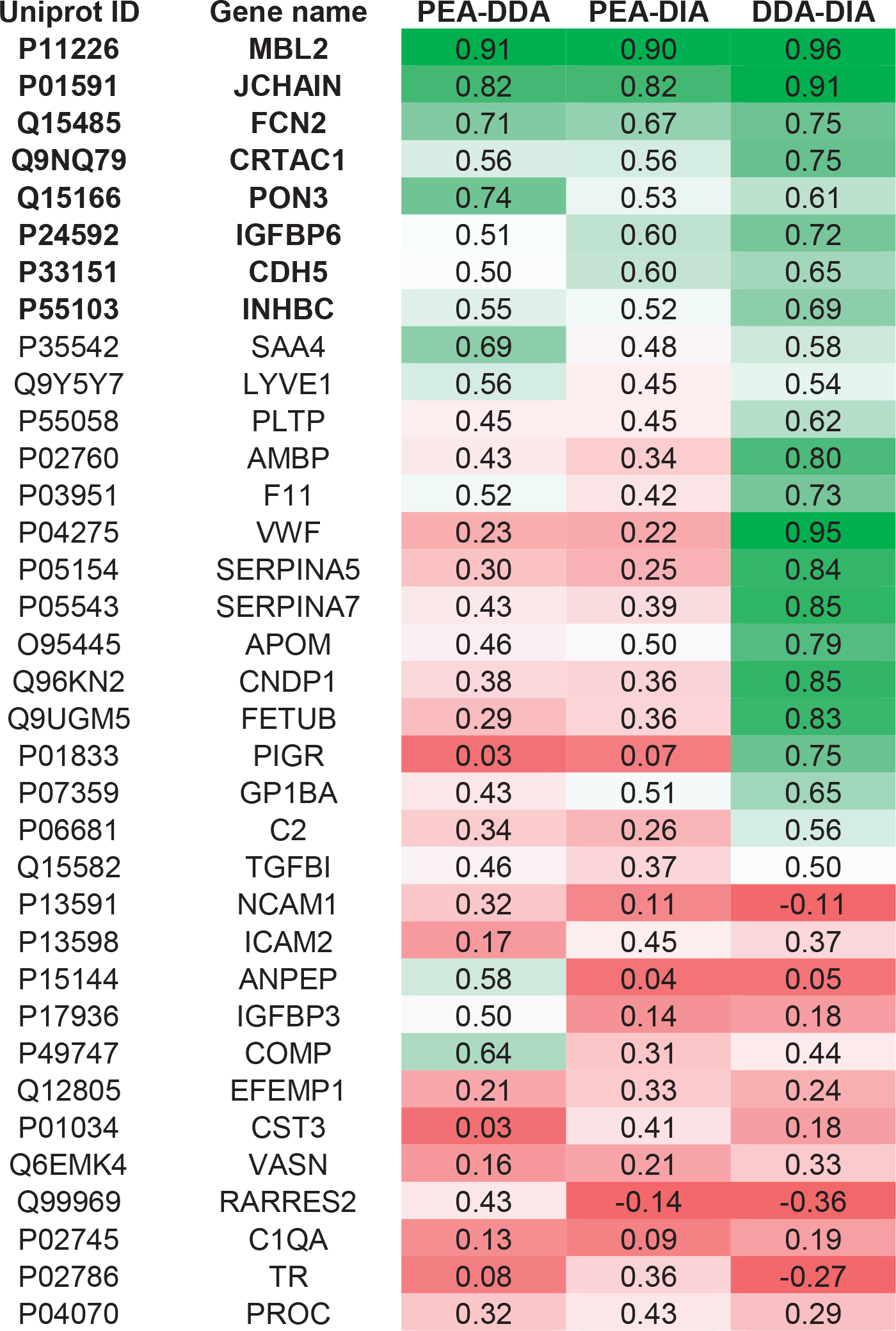
Inter-platforms correlation. Spearman coefficient R of the 35 overlapping proteins for each pairwise platform comparison. Proteins highlighted in bold show a moderate to a very high positive correlation (R > 0.5). The degree of positive (green) or negative (red) correlation is shown by the grade color scale heatmap.

| Uniprot ID | Gene name | PEA-DDA | PEA-DIA | DDA-DIA |
| --- | --- | --- | --- | --- |
| **P11226** | **MBL2** | 0.91 | 0.90 | 0.96 |
| **P01591** | **JCHAIN** | 0.82 | 0.82 | 0.91 |
| **Q15485** | **FCN2** | 0.71 | 0.67 | 0.75 |
| **Q9NQ79** | **CRTAC1** | 0.56 | 0.56 | 0.75 |
| **Q15166** | **PON3** | 0.74 | 0.53 | 0.61 |
| **P24592** | **IGFBP6** | 0.51 | 0.60 | 0.72 |
| **P33151** | **CDH5** | 0.50 | 0.60 | 0.65 |
| **P55103** | **INHBC** | 0.55 | 0.52 | 0.69 |
| P35542 | SAA4 | 0.69 | 0.48 | 0.58 |
| Q9Y5Y7 | LYVE1 | 0.56 | 0.45 | 0.54 |
| P55058 | PLTP | 0.45 | 0.45 | 0.62 |
| P02760 | AMBP | 0.43 | 0.34 | 0.80 |
| P03951 | F11 | 0.52 | 0.42 | 0.73 |
| P04275 | VWF | 0.23 | 0.22 | 0.95 |
| P05154 | SERPINA5 | 0.30 | 0.25 | 0.84 |
| P05543 | SERPINA7 | 0.43 | 0.39 | 0.85 |
| O95445 | APOM | 0.46 | 0.50 | 0.79 |
| Q96KN2 | CNDP1 | 0.38 | 0.36 | 0.85 |
| Q9UGM5 | FETUB | 0.29 | 0.36 | 0.83 |
| P01833 | PIGR | 0.03 | 0.07 | 0.75 |
| P07359 | GP1BA | 0.43 | 0.51 | 0.65 |
| P06681 | C2 | 0.34 | 0.26 | 0.56 |
| Q15582 | TGFBI | 0.46 | 0.37 | 0.50 |
| P13591 | NCAM1 | 0.32 | 0.11 | -0.11 |
| P13598 | ICAM2 | 0.17 | 0.45 | 0.37 |
| P15144 | ANPEP | 0.58 | 0.04 | 0.05 |
| P17936 | IGFBP3 | 0.50 | 0.14 | 0.18 |
| P49747 | COMP | 0.64 | 0.31 | 0.44 |
| Q12805 | EFEMP1 | 0.21 | 0.33 | 0.24 |
| P01034 | CST3 | 0.03 | 0.41 | 0.18 |
| Q6EMK4 | VASN | 0.16 | 0.21 | 0.33 |
| Q99969 | RARRES2 | 0.43 | -0.14 | -0.36 |
| P02745 | C1QA | 0.13 | 0.09 | 0.19 |
| P02786 | TR | 0.08 | 0.36 | -0.27 |
| P04070 | PROC | 0.32 | 0.43 | 0.29 |

**FIGURE 4.**
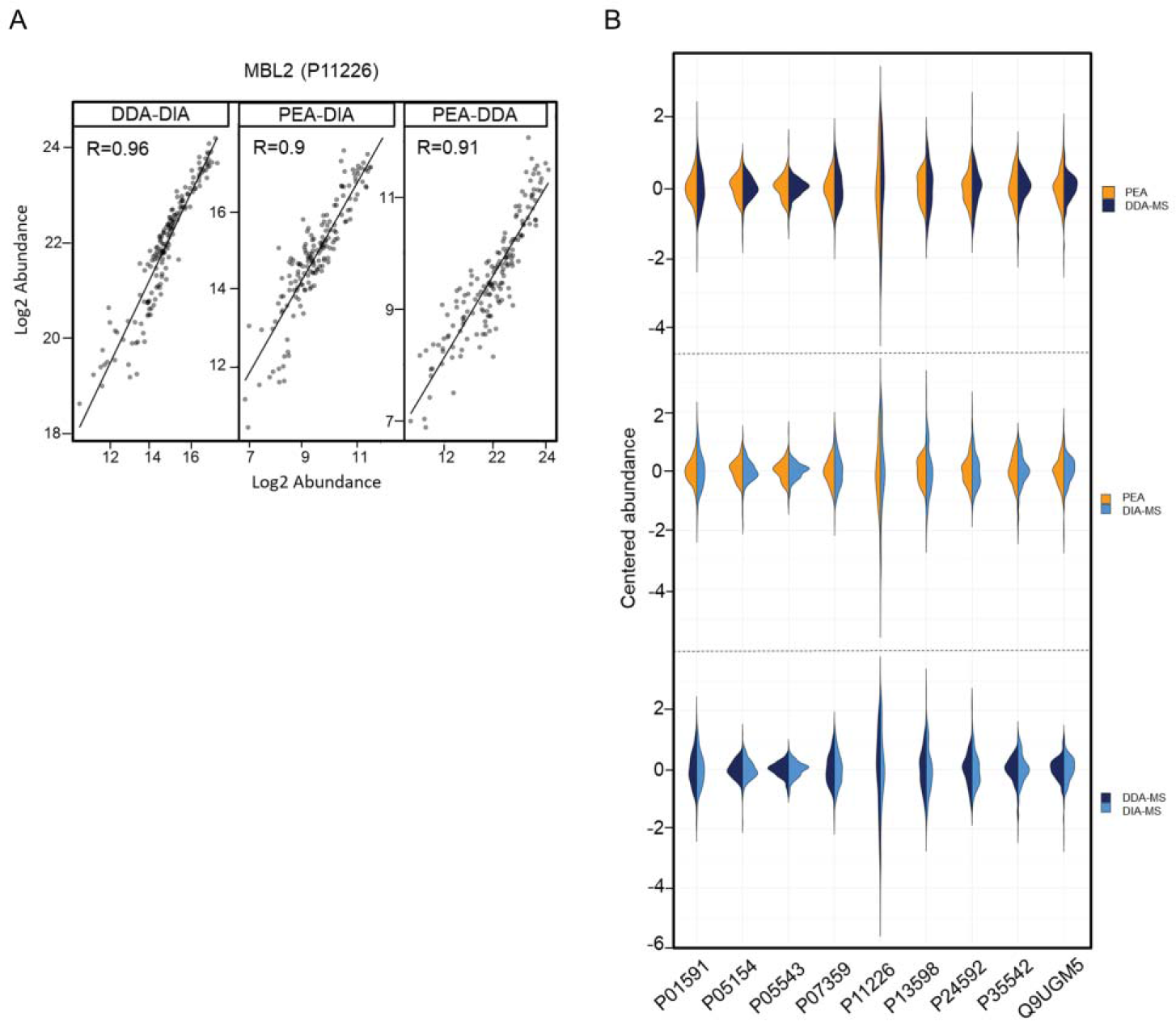
Spearman correlation of the 35 overlapping proteins. A. Plots showing the interplatform correlations for MBL2 (P11226) with respective Spearman correlation coefficient R. B. Violin plots of the z-score transformed centered data showing the distribution shape of nine not rejected proteins in all three platforms.

#### 2.3 Gender-specific correlation analysis of the overlapping proteins identified by PEA, DDA- and DIA-MS

Thirdly, statistical analysis was performed using gender, a key biological variable that contributes to individual variation. Given the comparable proportion of males and females in our cohort (males n=90 and females n=83), we carried out a gender-specific differential expression analysis. A Welch’s two-sample t-test was computed for all proteins within platform, followed by Benjamini-Hochberg (FDR) multiple testing correction at an alpha level of 0.05 (**Supplemental Table S9** and **Supplemental Figure S1**). We calculated 130 proteins significantly regulated in the PEA dataset and 68 proteins in the MS datasets, with 16 and 15 proteins uniquely present in the DDA and DIA-MS datasets, respectively. The Upset bar plot graph visualizes the overlap between the analytical platforms based on gender differentiation (Fig. 5A). Noteworthy is that, for the significant proteins that are also overlapping - IGFBP6 (Insulin-like growth factor-binding protein 6, P24592), F11 (Coagulation factor XI, P03951), PROC (Vitamin K-dependent protein C, P04070) and SERPINA7 (Thyroxine-binding globulin, P05543) – the direction of effect is the same. Although the effect size between platforms cannot be assumed to have exactly the same magnitude, the direction of effect is an important indicator of reproducibility of biological insight, i.e. the same significant up- or downregulation can be inferred from all platforms. Indeed, plotting the effect sizes of the overlapping proteins from each platform against each other (Fig. 5B) shows a clear similarity of direction in the effects (significant or not), indicating that biological outcomes are reproducible between platforms.

**FIGURE 5.**
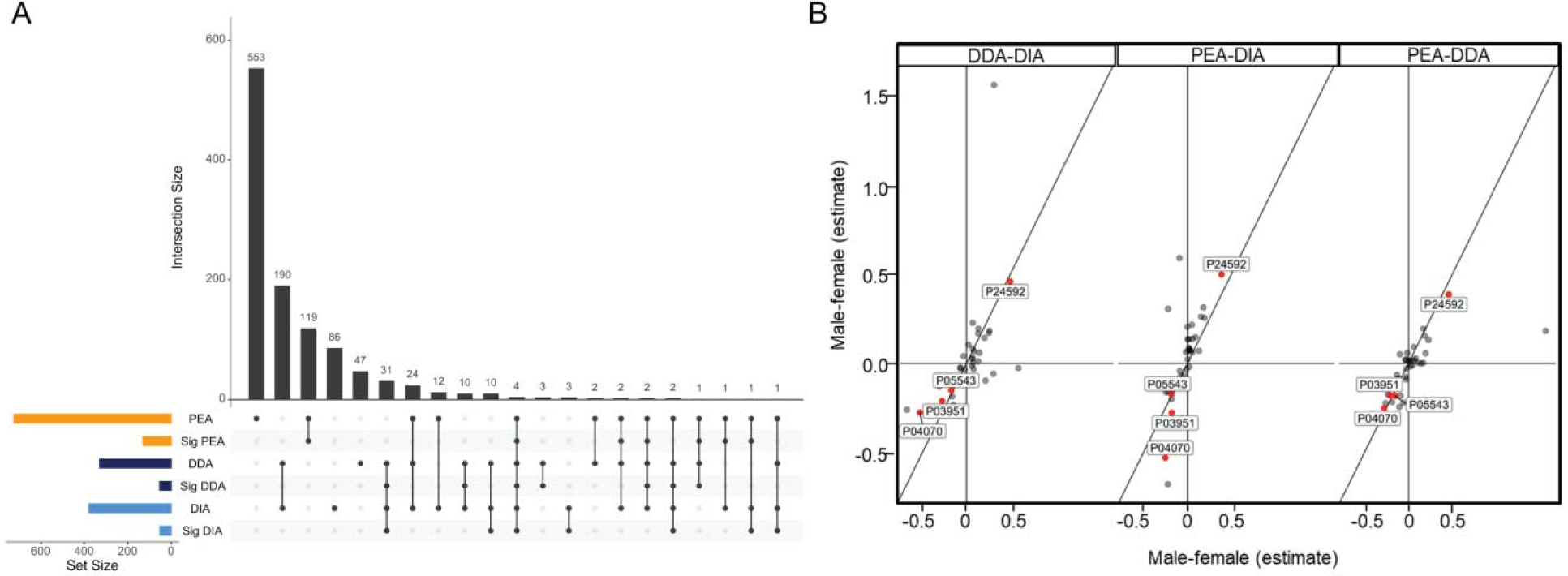
A. Upset plot summarizing the gender specific differential protein expression analysis for proteins measured by PEA and MS. Proteins were tested for significant differential expression between genders (log2 fold change expression) using Welch’s two-sample t-test followed by Benjamini-Hochberg FDR correction (*p* < 0.05). The horizontal bar graph at the bottom left shows for each method the total number of proteins identified in > 25% of the samples and the number of significantly expressed proteins (abbreviated Sig.). The vertical bar graph at the top quantitates the number of proteins shared between different sets of methods. The specific methods in each set are indicated with solid points below the bar chart. B. Correlation plot of the male-female estimate expression ratio of the 35 overlapping proteins (log2 fold change.) The four gender-related significantly expressed proteins are marked with a red dot.

## Discussion

We have investigated the plasma proteome of human samples derived from a population-based biobank by using two representative methods of targeted and untargeted proteomics, i.e. the Olink Proximity Extension Assay and LC-MS/MS in data-dependent and -independent acquisition modes. The final aim of this study is to show a new perspective into the application of currently available technologies in plasma proteomics. Unmet clinical needs for plasma biomarkers deal with the lack of tools with adequate sensitivity and specificity extended to a large dynamic range; reproducibility and multiplexing are also essential for the clinical translation. Both targeted and untargeted proteomics-based technologies have their own strengths and weaknesses, and often large deals of resources are invested to optimize and improve the performance of a single method. Instead, we propose here to combine two complementary methodologies to deeply explore the plasma proteome while keeping up reproducibility and sensitivity. Driven by the need to merge the strengths of untargeted LC-MS/MS plasma profiling and targeted PEA technology, we compared the performances of both methods by screening 173 human plasma samples. In terms of proteome coverage and detectability, we demonstrated that MS measurements in DDA and DIA modes largely identified identical proteins (276 IDs). DIA-MS outperformed DDA-MS workflow in terms of total number of identified proteins; however, the proteome coverage in more than 25% of the samples is comparable between DDA- and DIA-MS measurements. More than 90% of proteins included in the eight Olink panels were detected above the limit of detection in all samples, indicating an excellent detectability of the assays in human blood plasma from the general population. For all eight panels, we achieved an even higher protein detectability than the expected coverage measured by Olink on EDTA plasma from healthy donors (Validation Data Documents, https://www.olink.com/resources-support/document-download-center/). After confirming high stability and intra-assay reproducibility of each platform, we based our technical comparison on the proteins identified and quantified by all three methods. The small overlap between MS and PEA clearly shows that these technologies target two different fractions of the proteome, and they meet in the high abundance proteins region. We have analyzed the measurement reproducibility of 35 overlapping proteins in a pairwise statistical comparison. The data distribution of the measurements is highly reproducible across methods, especially between PEA and DDA-MS where the similarity of the distributions is even higher than between MS-acquisition modes. A deeper analysis of the data spread showed that the majority of the proteins have a narrow spread around the mean, which leads to a weak correlation of the measurements across methods. This concept is highlighted by the only protein, MBL2 (Mannose-binding protein C), which showed a broad data spread in all three measurements and thereby resulting in an excellent inter-assay correlation. The large variation in the MBL2 blood concentration is explained by six polymorphisms in the promoter 1 and exon 1 region of the MBL2 gene (27). These genetic variants significantly affect the serum MBL2 level, with severe consequences on the ability of MBL2 in fighting infections (28, 29). Our correlation analysis of MBL2 across several MS workflows and PEA suggests the robustness of this biomarker for future clinical applications. We finally introduced the biological variable gender to our statistical analysis to evaluate if similar conclusions can be drawn with regard to the overlapping proteins. We observed the same trend of gender related differential expression for the 35 proteins across platforms, including four proteins having statistically significant abundance ratio male-female in all three methods. This evidence takes on an important value when analytical platforms are combined to investigate biological features of a population-based cohort. Gender-specific plasma proteome holds considerable value in clinical practice, and gender differences in plasma biomarker levels have been widely reported in cancer, cardiovascular, neurological and pulmonary diseases (30–33). Our data suggest that we can merge the results of the PEA and MS platforms to reach greater proteome coverage and more biological insights. MS has proved since decades the ability to in-depth profiling a large amount of pathophysiological conditions at protein level, gaining attention and expectations in many branches of clinical proteomics. A quite unique strength of high-resolution MS/MS is the ability to map and quantify a large variety of post-translational modifications (PTMs) (34); PTM signatures are increasingly reported as potential biomarkers and more than 95% of current data on PTMs derives from MS-based proteomic studies (35). Another noteworthy advantage offered by one of the latest developed MS workflows, named middle-down proteomics, is the outstanding peptide sequence coverage (up to 95%) achieved by decreasing the peptide digestion complexity. This workflow will improve the identification of splice variants and protein isoforms, as well as co-occurring neighboring PTMs (36). These distinctive features make MS a key element in the study of the plasma proteome. Our results further support the high performance of MS to reliably identify and quantify more than 300 circulating proteins. In line with current findings in MS-based plasma proteome profiling, we mainly observe abundant functional proteins such as serum albumin, apolipoproteins, immunoglobulins, acute phase proteins and coagulation cascade components. This large fraction of high abundant proteins involved in the inflammation and lipid metabolism has enormous value for the profiling of cardiovascular and metabolic diseases (37, 38). However, we faced the limit of MS to dig into the low -abundant and -molecular weight fraction of the human plasma proteome. High-throughput multiplex immunoassays such as Olink PEA has made that hidden and hard-to-reach plasma proteome fraction enriched in biologically relevant small molecules accessible. The eight panels used in this study provided quantification of 25 cytokines, among them chemokines and interleukins, and five peptide hormones; none of these proteins was detected by MS. This illustrates the benefits of merging the comprehensiveness of MS and the sensitivity of PEA and the unprecedented potential resulting from their complementation. We have seen that the limitations of MS-based approaches are sort of strengths of PEA, and vice versa, which supports the complementarity between the two analytical technologies. Over the last decade we have faced the expanding growth of plasma proteomics methodologies driven by the urge to provide reliable diagnostic biomarkers. A key for a successful translation into clinics relies on screening multi-protein panels rather than a single protein, and both untargeted and targeted approaches can achieve this goal. Only through the complementation of diverse proteomics technologies, each with its own benefits and disadvantages, we will be able to explore the vast and rich world of the plasma proteome.

## Acknowledgements

The KORA study was initiated and financed by the Helmholtz Zentrum München – German Research Center for Environmental Health, which is funded by the German Federal Ministry of Education and Research (BMBF) and by the State of Bavaria. Furthermore, KORA research was supported within the Munich Center of Health Sciences (MC-Health), Ludwig-Maximilians-Universität München, as part of LMUinnovativ. We gratefully acknowledge Dr. Marianne Sandin for her valuable support and commitment. We are very thankful to Andrea Ballagi and Katarina Hörnaeus from Olink Uppsala for their cooperation and assistance running this study. We dedicate this work to the loving memory of our colleague and dear friend Fabian Metzger.

## Data Availability

Informed consents given by KORA study participants do not cover data posting in public databases. However, the KORA data is available given approval of online requests at the KORA Project Application Self-Service Tool (https://epi.helmholtz-muenchen.de/).

## Abbreviation

PEA: Proximity Extension Assay
DDA: Data-dependent acquisition
DIA: Data-independent acquisition
LC-MS/MS: liquid chromatography mass spectrometry
FDR: false discovery rate
ID: identification
KORA: Cooperative Health Research in the Region of Augsburg
Olink panels: C-MET cardiometabolic; CVDII cardiovascular II; CVDIII cardiovascular III; ONCII oncology II; ONCIII oncology III; DEV development; IMM immune response; NEU neurology

